# Mapping the influence of the gut microbiota on small molecules in the brain through mass spectrometry imaging

**DOI:** 10.1101/2020.03.13.987164

**Authors:** Heather Hulme, Lynsey M. Meikle, Nicole Strittmatter, John Swales, Gregory Hamm, Sheila L. Brown, Simon Milling, Andrew S. MacDonald, Richard J.A. Goodwin, Richard Burchmore, Daniel M. Wall

**Author notes:** Corresponding author: Corresponding author address: Dr. Daniel (Dónal) M. Wall, Institute of Infection, Immunity and Inflammation, College of Medical, Veterinary and Life Sciences, Sir Graeme Davies Building, University of Glasgow, 120 University Place, Glasgow G12 8TA.

## Abstract

**Background:** The gut microbiota is known to influence virtually all facets of human health. Recent work has highlighted a potential role for the gut microbiota in neurological health through the microbiome-gut-brain axis. Microbes can influence the brain both directly and indirectly; through neurotransmitter production, induction of host immunomodulators, or through the release or induction of other microbial or host molecules.

**Methods:** Here we used mass spectrometry imaging (MSI), a label-free imaging tool, to map the molecular changes that occur in the murine gut and brain in germ-free, antibiotic-treated and control mice.

**Results:** We determined the spatial distribution and relative quantification of neurotransmitters and their precursors across brain and gut sections in response to the microbiome. Using untargeted MSI of small molecules, we detected a significant change in the levels of four identified metabolites in the brains of germ-free animals compared to controls; vitamin B5, 3-hydroxy-3-methylglutaric acid, 3-methyl-4-(trimethylammonio)butanoate and 4-(trimethylammonio)pentanoate. However, antibiotic treatment induced no significant changes in these metabolites in the brain after one week of treatment.

**Conclusions:** This work exemplifies the utility of MSI as a tool in determining the spatial distribution and quantification of bacterial and host metabolites in the gut and brain whilst also offering the potential for discovery of novel mediators of microbiome-gut-brain axis communication.

## Background

Deciphering the complex bi-directional communication across the microbiome gut brain (MGB) axis remains a challenging prospect. The composition and stability of the gut microbiome is now proposed to be a significant contributor to human health with changes in its composition suggested as a contributing factor in a number of neurological conditions. Diverse human neurological disorders, ranging from autism spectrum disorders (ASDs) and attention-deficit/hyperactivity disorder (ADHD) to Alzheimer’s and Parkinson’s disease, have linked gastrointestinal abnormalities or changes in the gut microbiome (*1*–*6*). Similarly, altered levels of certain bacterial species in the gut have been linked with depression (*7*, *8*). Despite the deficiencies of germ free (GF) animal models, their use for investigating links between the gut microbiota and the brain have proved informative and are helping to uncover the influence of bacteria on neurotransmitter levels alongside other bacterial and host metabolites. Their neuro-developmental abnormalities including increased blood-brain-barrier (BBB) permeability, alterations in abundance and maturity of microglia cells, and reduced myelination (*9*–*11*) have been well documented but their use, alongside antibiotic treated (ABX) mice, have provided further evidence that supports the importance of a stable and healthy gut microbiota in maintaining normal cognitive function and development (*12*). GF mice display reduced anxiety-like behaviour and an increased response to stress that is fully alleviated upon colonisation with *Bifidobacterium infantis* (*12*–*15*) while murine ABX treatment models reinforce the significance of the MGB axis with treated mice showing impaired cognition and significantly altered behaviours that can be linked to absence of particular bacteria (*16*–*19*). These effects are likely mediated by the numerous bacterial molecules including short chain fatty acids (SCFAs), phenolic acids, quorum sensing peptides, peptidoglycan and lactic acid, that have been proposed to exert influence across the MGB axis (*20*–*22*). SCFAs both cross and modulate the permeability of the BBB, while animals and humans are also dependent on intestinal microbes to produce or supplement their vitamin needs (*10*, *23*, *24*).

Signalling across the MGB axis can occur via the vagus nerve, bacterial modulation of host immune responses or host neurotransmitter production, and signalling through bacterial neurotransmitters or other microbial molecules. Microbes produce a number of neurotransmitters including gamma aminobutyric acid (GABA), norepinephrine and dopamine, serotonin, and acetylcholine while bacterial production or depletion of essential neurotransmitter intermediates such as tryptophan can also affect host production (*14*, *19*, *25*–*30*). Manipulation of the microbiome can also alter host neurotransmitter levels as microbes stimulate their intestinal production with depletion of the microbiome, or its supplementation with probiotic bacteria, capable of altering specific levels of neurotransmitters even in the brain (*31*–*35*).

Understanding how the gut microbiota influence the brain, and the complex network of molecules and neurotransmitters that mediate this influence, is a significant challenge requiring novel tools and approaches. Mass spectrometry imaging (MSI) is a molecular imaging tool that can be applied to understand and map biological systems with recent successes in mapping microbial interactions with their environment (*36*–*38*). MSI maps the distribution of small molecules across a tissue section independent of any label, so no prior knowledge of the molecules present is required. The ability to detect spatial distribution and abundance of thousands of compounds simultaneously in tissue sections makes this a powerful approach for studying the MGB axis. Here, using MSI, we detected significant differences in neurotransmitter levels and those of their precursors in the guts of both GF and ABX mice. However in the brain no differences were detected in neurotransmitter levels in GF mice compared to controls, while only tryptophan levels significantly changed in the brains of ABX mice. Through untargeted MSI we also discovered 3 metabolites were significantly changed in the GF brain. Two of these were identified as 3-hydroxy-3-methylglutaric acid (3-HMG) and pantothenic acid (vitamin B5). 3-HMG, a metabolite associated with oxidative stress, was significantly increased in the GF brain, while vitamin B5, which can be produced by the microbiome, was present at a lower level in GF brain and is implicated in brain health.

This work indicates that the brain remains largely protected from microbiome changes in the gut and indicates plasticity, not just in the brain but across the MGB axis in GF mice.

## Methods

### Animal models used in this study

#### Germ-free studies

All germ-free (GF) work was undertaken at the University of Manchester Gnotobiotic Facility. Five GF and specific pathogen free (SPF) mice were used in this study, which were male mice, aged 7-8 weeks, on a C57BL/6J background strain. Both experimental groups were fed the same pelleted diet that was irradiated with 50 kGy to ensure sterility. The Manchester Gnotobiotic Facility was established with the support of the Wellcome Trust (097820/Z/11/B), using founder mice obtained from the Clean Mouse Facility (CMF), University of Bern, Bern, Switzerland.

#### Antibiotic studies

All work involving antibiotic (ABX) treatment was undertaken at the University of Glasgow. Five ABX treated animals and corresponding untreated controls were used in these studies and were male mice, aged 7-8 weeks, on a C57BL/6J background strain. The ABX cocktail consisted of 1 mg/ml gentamicin, 1 mg/ml neomycin and 0.5 mg/ml vancomycin in sterile distilled drinking. ABX supplemented drinking water was provided ad libitum for a period of 1 week and refreshed every 2 days. The untreated controls were given sterile drinking water without ABXs ad libitum, which again was refreshed every two days. Both treated and control groups were fed the same standard chow. All antibiotics selected for this study were chosen based on their specificity for the gut microbiota; to the best of our knowledge the antibiotics used are not absorbed by the intestine. Approval for these procedures was given prior to their initiation by internal University of Manchester and University of Glasgow ethics committees and by the U.K. Home Office under licenses 70/7815, PPL40/4500, P64BCA712 and P78DD6240.

### Tissue processing

Mice were culled by cervical dislocation and brains and colons were removed. Brains were placed in a mould and were immediately frozen using a slurry of dry-ice and ethanol to maintain structural integrity and to ensure that all biochemical processes were halted. The colon samples were then cut lengthwise over ice and the faecal matter was removed and the remaining GI tissue was then rolled using the ‘Swiss roll technique’ before embedding in 2.5% medium viscosity carboxymethyl cellulose (Sigma-Aldrich, Dorset, UK). Both brains and colons were sectioned using a CM3050S cryostat microtome (Leica Biosystems, Nussloch, Germany) to 10 μm thickness at −18°C. Sagittal brain sections were either thaw mounted onto indium tin oxide (ITO) coated slides for Matrix Assisted Laser Desorption/Ionization (MALDI)-MSI or normal microscope slides for desorption electroSpray ionisation (DESI)-MSI. Stereotactically matched sections were selected to ensure that all comparisons were performed using corresponding brain regions. Consecutive sections to those for MSI were collected for histology purposes. Slides were prepared and stored at −80°C until required for analysis. Prior to derivatisation, matrix application or analysis, the slide was taken from −80°C and brought to room temperature under a stream of air.

### Neurotransmitter derivatisation

The derivatisation of primary amine neurotransmitters was performed as previously described (*39*). Nine milligrams of 2,4-diphenyl-pyranylium tetrafluoroborate (DPP-TFB) was added to 1.2 mL of 100% methanol and sonicated for 20 minutes (min). This was then gradually added to 6 mL of 70% methanol in water with 3.5 μL of trimethylamine. This solution was sprayed onto the tissue for derivatisation using an automated sprayer (TM-Sprayer, HTX Technologies); 30 passes were performed using a nozzle temperature of 75°C, velocity of 1100 mm/min, flow rate of 80 μL/min, and gas pressure of 6psi. After coating, the slide was incubated in a petri dish with vapour from a 50 % methanol/water solution 3 times for 5 minutes each.

### DESI-MSI analysis

DESI-MSI was performed on a Q-Exactive mass spectrometer (Thermo Scientific, Waltham, MA, US) equipped with an automated 2D DESI source (Prosolia Inc, Indianapolis, IN, USA). A home-built DESI sprayer assembly, as described previously (*40*), was used with the spray tip positioned at 1.5 mm above the sample surface and at an angle of 75°. The distance between the sprayer to mass spectrometer inlet was 7 mm with a collection angle of 10° and <<1mm distance between inlet and sample surface. The spray solvent was methanol/water (95:5 *v/v*), delivered at 1.5 μL/min using a Dionex Ultimate 3000 pump (Thermo Scientific, Waltham, MA, US) at a spray voltage of ±4.5 kV. Nitrogen was used as the nebulisation gas at a pressure of 7 bars. General instrument settings used to image specific molecules are shown in Table 1. For acquisition of MS/MS spectra, an injection time of 300 ms, mass resolution of 70000 and a mass isolation window of ± 0.3 Da was used. For MS/MS imaging of metabolites with mass to charge (*m/z*) ratios 161.044 and 218.103, various fragmentation HCD settings were used, at a spatial resolution of 100 μm.

Data was recorded as individual line scans and converted into imzML format using imzML converter version 1.1.4.5 (*41*) and visualised using MSiReader version 0.09 (*42*). All intensities shown are raw ion intensities and 1^st^ order linear interpolation was used for image generation. All mean intensities of the molecules of interest were determined across the entire tissue section. The data were transformed to log10 and a Shapiro-Wilk normality test was performed on the data to check for a normal distribution. If the data passed the normality test an unpaired t-test was performed. If the data failed the test a Mann-Whitney U test was performed. When biological replicates were analysed over two separate DESI-MSI experiments a paired t-test was performed instead.

**Table 1.**
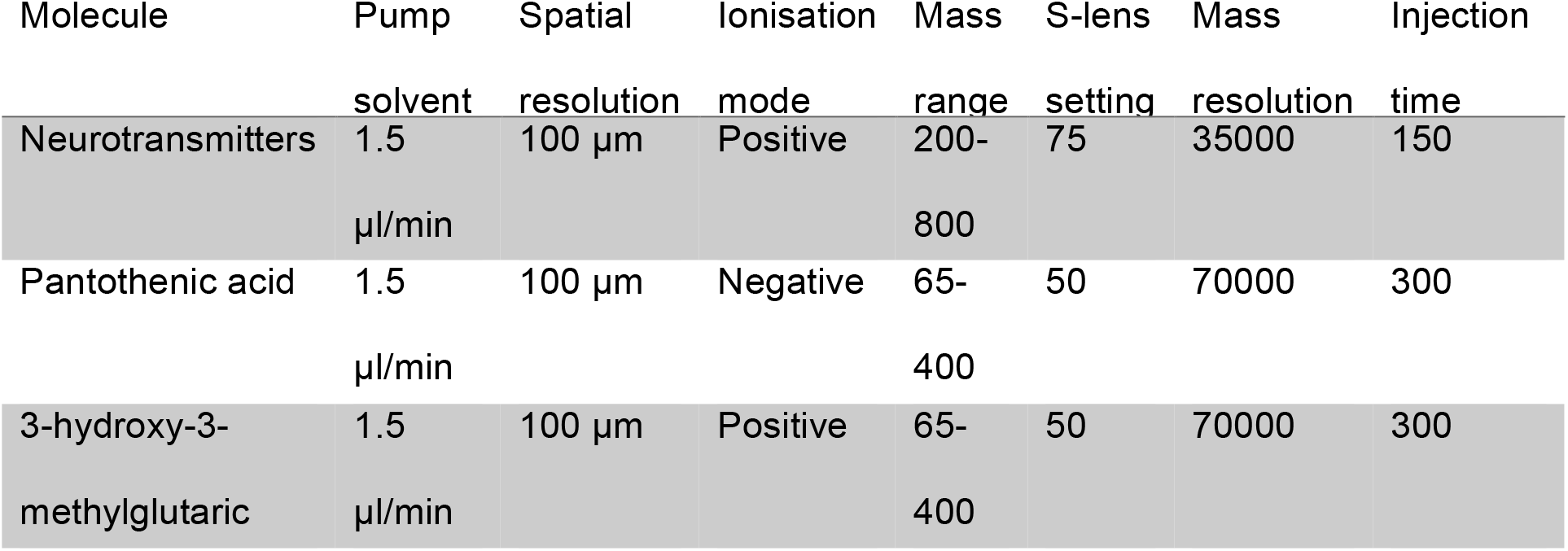
DESI-MSI parameters

### H&E staining of brains and colons

Brain and colon sections that had undergone MSI analysis were H and E stained post-imaging to permit localisation of candidate metabolites and neurotransmitters to specific brain regions. Sections were fixed on the slide in ice cold 75% acetone and 25% ethanol for 10 min, and air dry for a further 10 min. Slides were placed in water for 2 min, submerged in haematoxylin (Sigma-Aldrich, Poole, Dorset, UK) for 2 min and immediately rinsed in cold running water. The slides were then dipped for 3 sec in acid alcohol 0.5% (Atom Scientific Hyde, Cheshire, UK), and rinsed in water before submerging in Scott’s tap water (Atom Scientific Hyde, Cheshire, UK) for a further 30 sec. The sections were counter-stained with eosin (Sigma Aldrich, Poole, Dorset, UK) for 2 min and washed in water. Sections were then dehydrated in increasing concentrations of ethanol (70 % ethanol for 30 sec, 90 % for 1 min, and twice for 3 min in 100 % ethanol), cleared in xylene (twice for 3 min), and cover slipped using DPX mounting media (Atom Scientific, Hyde, Cheshire, UK).

## Results

### Targeted neurotransmitters remain unaffected by the lack of a gut microbiota

The brains and colons from germ free (GF) and conventionally colonized, specific pathogen free (SPF) control mice were first compared by MSI using a targeted approach. DPP-derivatisation of primary amines was performed to allow targeted imaging of neurotransmitters, neurotransmitter precursors and neurotransmitter metabolites; serotonin, tryptophan, dopamine, tyrosine, 3-methoxy-tyramine, GABA and glutamate (Figure 1) (*39*, *43*, *44*). Significant differences in the levels of several neurotransmitters were observed in the gut. Serotonin was lower (3.8-fold decrease) in the GF mouse gut (Fig. 1b; *P* ≤0.01), whilst glutamate (1.3-fold increase) (Fig. 1b; *P* ≤0.05) and the dopamine precursor tyrosine (1.8-fold increase) (Fig. 1b; *P* ≤0.01) were significantly increased in the GF gut. However, these changes were not reflected in the corresponding brain sections of GF mice where no significant differences were identified for any of the seven neurotransmitters or their precursors (Fig. 1a). There were also no differences detected in the levels of tryptophan, dopamine, 3-methoxy-tyramine or GABA in either the brain or gut sections from GF and SPF mice (Supplementary Figure 1).

**Figure 1.**
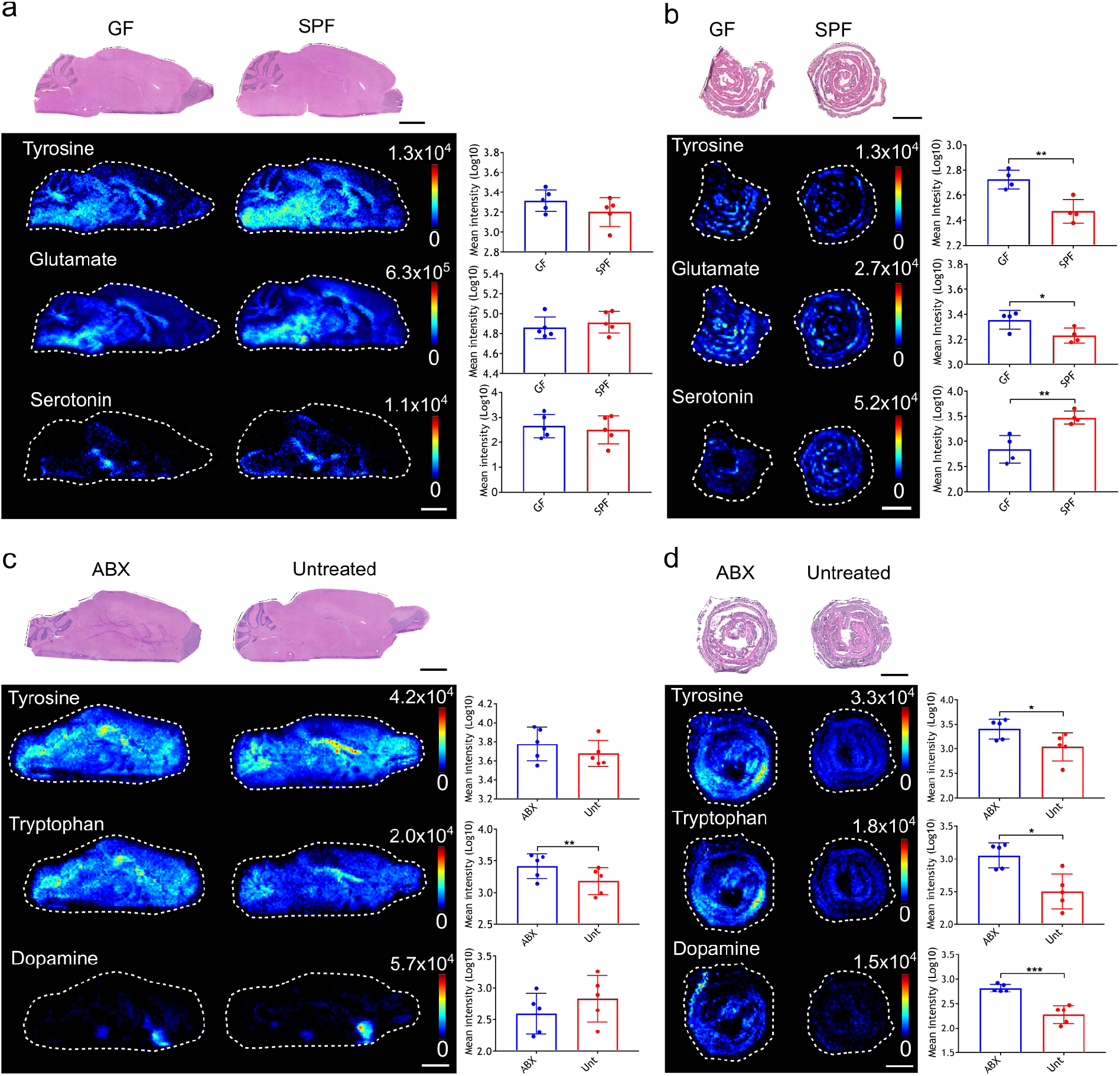
Effects of the gut microbiome on neurotransmitters, neurotransmitter precursors and neurotransmitter metabolites in the murine brain and gut. DESI-MS images of various DPP-derivatised neurotransmitters, neurotransmitter precursors or neurotransmitter metabolites in: (a) germ-free (GF) and specific-pathogen-free (SPF) mouse brains (N=5), (b) GF and SPF mice colons (N=4), (c) antibiotic treated (ABX) and untreated (Unt) mouse brains (N=5), (d) and ABX and Unt mice colons (N=5). H and E stained sections are shown, which are the same tissue sections that underwent DESI-MSI analysis. The bar plots show the average metabolite abundance (ion intensity) across the whole tissue sections in the different groups tested. For (b), statistical analysis was performed using an unpaired t test. For (c) and (d), statistical analysis was performed using a paired t test, an unpaired t test or a Mann-Whitney U test where appropriate. Error bars represent standard deviation. *, p≤0.05; **, p≤0.01; ***, p≤0.001. Scale bars = 2 mm.

### MSI of neurotransmitters in the brains and colons of ABX treated mice

To determine whether an acute disruption of the microbiota over 7 days could have an influence on the levels or the localisation of the same neurotransmitters in the brain, the colon and the brains of ABX treated mice were imaged in a targeted manner.

Tyrosine levels were changed in ABX treated mice in a manner similar to that seen in GF mice with a significant increase in the colon (2.2-fold), but again this increase was not reflected in tyrosine levels in the brain of ABX treated animals (Fig. 1c and 1d; *P* ≤0.05). Unlike GF mice, ABX treated mice showed no change in the levels of serotonin in either the colon or brain, compared to untreated mice (Fig. 1c and 1d). However, there was significantly higher abundance of tryptophan in both the colon (3.2-fold) (*P* ≤0.05) and brain (1.7-fold) (*P* ≤0.01) of ABX treated mice compared to controls (Fig. 1c and 1d). This was noted across the whole brain. Dopamine levels were also significantly higher (3.3-fold) in the colons from ABX treated mice compared to untreated, but no significant change was detected in the brain (Fig. 1d; *P* ≤0.001). No significant difference was detected in levels of 3-methoxytyramine, glutamate or GABA in either the gut or brain sections imaged (Supplementary Figure 2).

### Untargeted MSI to detect novel molecular changes in GF mouse brains

As no significant changes were seen in several neurotransmitters in GF brains, untargeted imaging was performed using DESI-MSI on the colon and brains of GF mice to probe the MGB axis for molecular changes induced by microbiota disruption. Full scan spectra were collected from *m/z* 65-400 and *m/z* 250-1000, in both positive and negative ionisation mode, allowing detection of a wide range of metabolites and not targeted towards a particular group. Significant differences were detected in only three masses when comparing GF and SPF mouse brains. Two of these identified metabolites at *m/z* of 218.1030 and *m/z* 161.0446, both detected in negative ion mode, were selected for further analysis as putative identities could be assigned from online databases, as discussed further below. The third mass at *m/z* 160.133 was below the limit of detection in GF brains and could not be assigned an identity from online databases. Determination of the identity of this mass, the associated structures and bacterial origin has been recently described (Hulme *et al*., In press).

### Levels of 3-hydroxy-3-methylglutaric acid (3-HMG –m/z 161.0446) are increased in the GF brain

The molecule detected at *m/z* 161.0446 in negative ionisation mode, was found at significantly higher levels (*P* ≤0.0001) across the whole brain section of GF compared to SPF mice, with levels particularly high in the cerebellum (Fig. 2a). The change in the brain was not reflected in the gut as levels in the colon did not significantly change between GF and SPF control mice (Fig. 2b). The metabolite was identified as [M-H]- of 3-hydroxy-3-methylglutaric acid (3-HMG) through searching the Human Metabolome Database (*45*, *46*) and then confirmed using tandem mass spectrometry (MS/MS) analysis and comparison to an HMG standard (Supplementary figure 3). No significant difference in abundance of HMG was found in brains and colon after ABX treatment compared to controls (Fig. 2c and 2d).

**Figure 2.**
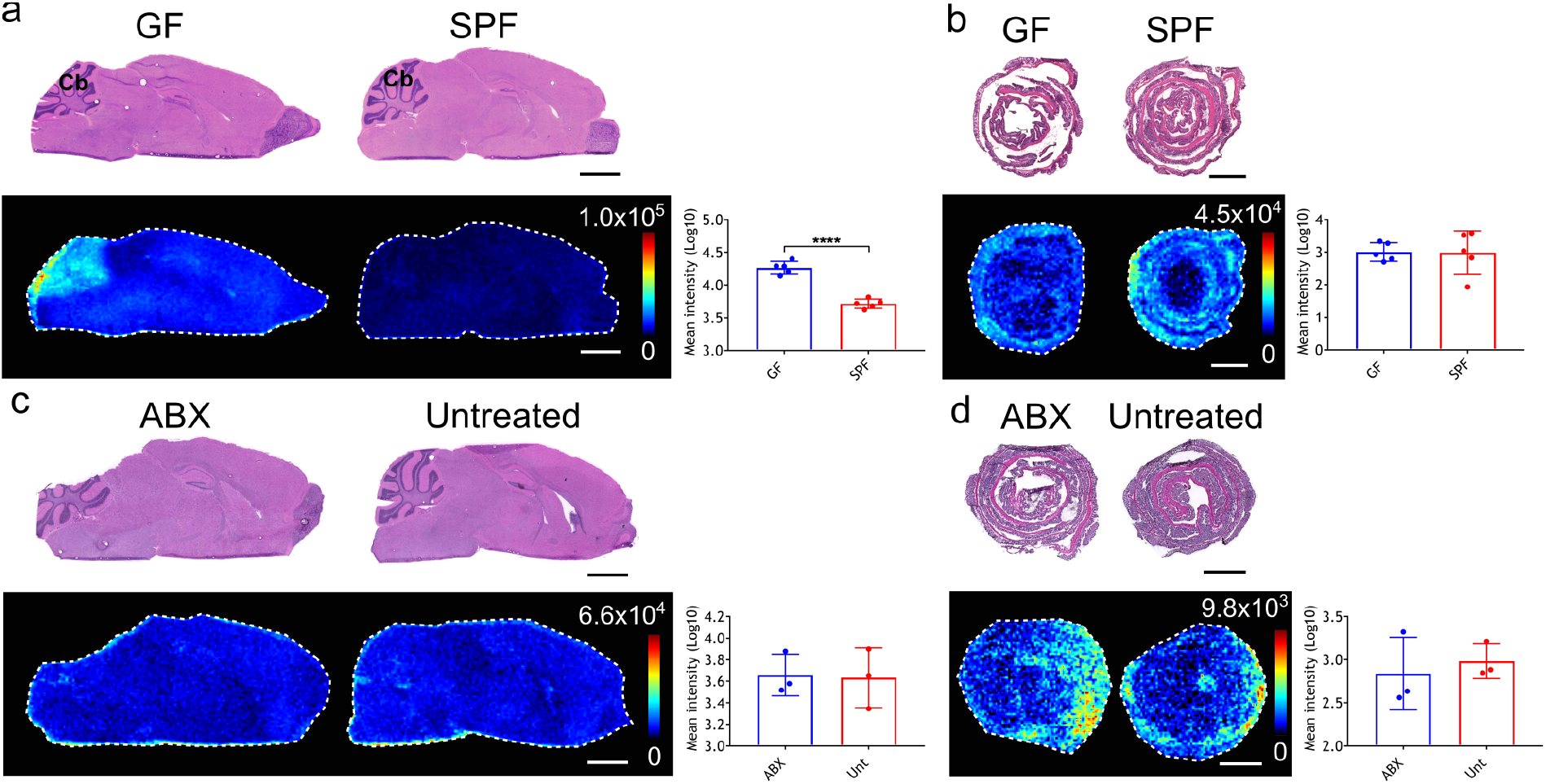
Impact of the gut microbiome on 3-hydroxy-3-methylgularic acid levels in the gut and brain. DESI-MS images of HMG in the (a) GF and SPF mouse brains (N=5), (b) GF and SPF mice colons (N=5), (c) ABX treated and Unt mouse brain (N=3), (d) and ABX treated and Unt colons (N=3). H and E stained sections are shown, which are the same tissue sections that underwent DESI-MSI analysis. The bar plots show the average HMG abundance (ion intensity) across the whole tissue sections in the different groups tested. Annotation in (a) Cb, cerebellum. Statistical analysis was performed using an unpaired t test. Error bars represent standard deviation N=5. ****, p≤0.0001. Scale bars=2 mm.

Quantification analysis of HMG revealed an average concentration of 0.56 μg/g of tissue in the GF brain compared to <0.01 μg/g of tissue in the SPF brain (Supplementary figure 4).

The concentration of HMG was particularly high in the cerebellum in the GF mouse brain at a concentration of 5.02 μg/g of tissue compared to <0.01 μg/g of tissue in the SPF mouse brains.

### Vitamin B5 (m/z 218.1030) levels are decreased in the brain of GF mice

The second of the unknown molecules had an *m/z* of 218.1030 in negative ionisation mode and was found to be significantly lower in the brain of GF mice compared to SPF controls (Fig. 3a; *P* ≤0.05). This difference was most obvious in the cerebellum, the hippocampus and the hypothalamus, where the molecule was most abundant in the SPF mice. This molecule was unchanged between GF and SPF colons. The metabolite was identified as [M-H]- of pantothenic acid, also known as vitamin B5, through database search and subsequent confirmatory MS/MS analysis.

**Figure 3.**
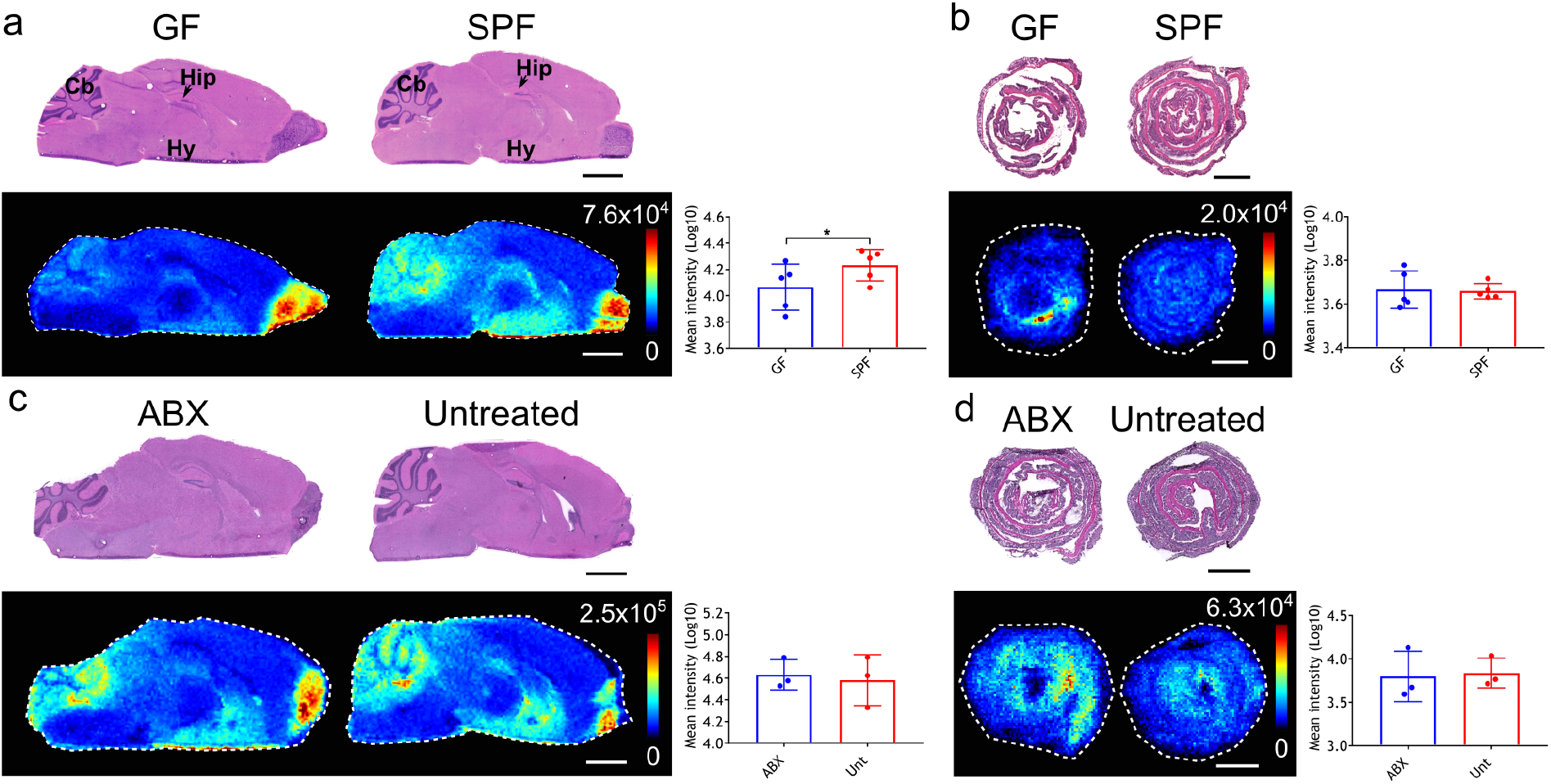
Effects of the gut microbiome on pantothenic acid (B5) levels in the gut and brain. DESI-MS images of B5 in the (a) GF and SPF mouse brains (N=5), (b) GF and SPF mice colons (N=5), (c) ABX treated and Unt mouse brain (N=3), (d) and ABX treated and Unt colons (N=3). H and E stained sections are shown, which are the same tissue sections that underwent DESI-MSI analysis. The bar plots show the average B5 abundance (ion intensity) across the whole tissue sections in the different groups tested. Annotation in (a) Cb, cerebellum; Hip, hippocampus; Hy, hypothalamus. Statistical analysis was performed using a paired t test. Error bars represent standard deviation N=5. *, p≤0.05. Scale bars=2 mm.

No difference was found in the levels of vitamin B5 in the brain or colon in ABX treated mice compared to untreated control mice (Fig. 3a and 3b).

Quantification analysis of vitamin B5 revealed an average concentration of 0.25 μg/g of tissue in the GF brain compared to 0.36 μg/g of tissue in the SPF brain.

## Discussion

Studying the MGB axis has significant potential to help us understand and potentially treat, via the microbiome, certain neurological conditions. However in order to achieve this goal a greater understanding of MGB communication is required. The techniques applied to date to discover mediators of communication across the MGB axis have typically been targeted toward specific neurotransmitters, metabolites or regions of the brain (*14*, *19*, *26*). Such approaches are often dependent on analyte extraction from brain tissue prior to analysis, meaning data regarding spatial localisation within the brain was limited. Here we applied MSI to study the metabolic processes along the MGB axis, permitting imaging of multiple neurotransmitters in the presence and absence of a microbiome while untargeted MSI also allowed the discovery of novel metabolites involved in MGB axis communication.

Serotonin, tryptophan, dopamine, tyrosine, 3-methoxytyramine, GABA and glutamate were first imaged to detect changes across the brain and gut of GF animals. No changes were detected in the brain whilst in the intestine only tyrosine and glutamate were increased and serotonin decreased in GF animals compared to SPF controls. Significantly lower levels of serotonin have previously been detected in the colon of GF mice along with increased levels of the serotonin precursor tryptophan (*32*). Whilst there was a trend towards increased levels of tryptophan in the present study this was not significant. No difference in serotonin or tryptophan levels were observed across the brain sections of GF mice in comparison to SPF animals here, in contrast to previous reported GF studies (*14*, *47*). However, both rat and mouse models have been used in these studies, complicating interpretation of data that has also indicated that these differences can be sex-specific. Whilst microbiome changes within mouse colonies as well as between mouse strains are well documented, phenotypic comparisons between GF rodents which lack a microbiome have been limited (*48*–*50*). One week antibiotic (ABX) treated mice were also tested for changes in levels of serotonin and tryptophan in the gut and brain compared to untreated controls. Tryptophan abundance increased in both the colon and the brain of ABX treated mice compared to untreated controls, but serotonin levels remained unchanged. This data mirrors that previously obtained with a similar ABX model, where ABX treatment increased systemic tryptophan whilst serotonin levels remained unaffected (*19*, *34*).

Dopamine and its precursor tyrosine were also affected by microbiota disruption or absence. Increased tyrosine and dopamine levels were detected in the colon in ABX treated mice while GF mice had increased intestinal tyrosine levels compared to controls. No difference was found in the abundance of either of these molecules in brains from either ABX treated or GF mice when compared to controls. Previous work found a decrease in the level of dopamine in the guts of GF mice, whilst another found no difference in the levels of dopamine in the colon after ABX treatment, although antibiotic treatments varied by constitution and duration between ours and previous studies (*28*)(*16*). As tyrosine can be metabolized by certain bacterial species, it is possible that reduction in these groups could lead to higher production of dopamine (*51*). Conversely the release of a biologically active, free form of dopamine in the gut via bacterial β-glucuronidase mediated breakdown of a conjugated form of dopamine could also play a significant role (*28*). Additionally, although there are no differences in the levels of dopamine in the brain, increased levels in the colon could have localised effects. There are dopamine receptors present in the intestine and dopamine, in a similar manner to serotonin, has been shown to increase water absorption from the gut and regulation of muscle contraction (*28*, *52*, *53*).

The neurotransmitter, and GABA precursor, glutamate was increased in the colon of GF mice compared to controls. Glutamate and GABA are the main excitatory and inhibitory neurotransmitters of the central nervous system, respectively (*54*). The increase in glutamate levels in GF mice was surprising given the number of bacteria known to produce glutamate in the intestine (*55*). This build-up of glutamate is not due to reduced conversion to GABA as we detected no corresponding difference in GABA levels (*56*). No increase in glutamate in the GF brain was detected in these GF mice with high intestinal glutamate levels, but glutamate is primarily metabolised in the splanchnic area and little enters into circulation from the gastrointestinal tract, instead exerting its significant localised effects on the gut including through stimulation of the vagus nerve (*57*, *58*).

Untargeted MSI indicated that three metabolites were significantly altered in the brain, vitamin B5 or pantothenic acid (*m/z* 218.1030) and 3-hydroxy-3-methylglutaric acid (3-HMG)(*m/z* 161.0446) and *m/z* 160.133 (Hulme et al., In press). The latter *m/z* was determined to be in fact a mixture of two molecules which are discussed in detail elsewhere (Hulme *et al*., In press). Vitamin B5 was significantly decreased in GF mice compared to SPF, whereas 3-HMG was increased. Further analysis of vitamin B5 and HMG determined that they were not significantly altered in ABX treated mice. Vitamin B5 is a precursor for co-enzyme A, which is an important molecule in many metabolic pathways including neurotransmitter production. Previously it was thought that only small amounts of vitamin B5 could be produced by the microbiota but recent work identified a previously uncharacterised group of *Clostridia* that harbour genes for pantothenic acid biosynthesis (*59*, *60*). Vitamin B5 is involved in oxidative metabolism and the synthesis of neurotransmitters including acetylcholine (*61*). Neurological symptoms of vitamin B5 deficiency were determined in early studies by inducing deficiency in human subjects, which resulted in defects in neuromuscular function and deterioration of mood (*62*). It has also recently been determined that levels of vitamin B5 producing bacteria change according to gestational age in preterm infants (*60*). The lower levels in the GF brain were particularly apparent in the cerebellum, hippocampus and hypothalamus brain regions. Despite the potential for a microbial origin for vitamin B5, no change was found in the levels of vitamin B5 in the colons of GF mice making it unclear what role, if any, the microbiota may play in the significant decrease in vitamin B5 levels in the brain. While vitamin B5 crosses into the brain through the BBB via a saturable transporter and levels are maintained in the brain at around 50 times the concentration found in the plasma, the mechanism of efflux of this vitamin from the brain is unclear and it may also be possible that increased BBB permeability in GF mice could lead to increased efflux from the brain (*63*, *64*)(*10*).

HMG, a metabolite involved in leucine degradation and ketogenesis was present at higher levels across the brains of GF mice compared to normal brains, but particularly in the cerebellum. Individuals with the genetic disorder 3-hydroxy-3-methylglutaric aciduria, which leads to a build-up of metabolites including HMG, suffer from neurological symptoms including seizures and abnormalities in the brain (*65*). It has been proposed that this accumulation of metabolites is directly affecting the brain with HMG inducing oxidative damage (*65*–*68*). Presence of HMG in the serum has been associated with increased gut permeability in children with environmental enteric dysfunction (*69*). Studies have shown that bacteria play a role in maintaining intestinal barrier function, therefore, the increased intestinal permeability seen in GF guts could be leading to higher levels of HMG in circulation compared to normal mice (*70*). Furthermore, intraperitoneal injections of HMG into rats leads to high levels accumulating in the brain of 7 day old rats but not in 30 day old rats in what was speculated to be a BBB permeability related effect (*65*). Therefore given the known BBB, and likely intestinal, permeability defects in GF mice, HMG, and potentially other metabolites, are likely entering the circulation and subsequently the brain at higher levels (*10*). Furthermore, no significant difference was found in the levels of HMG in the brain of ABX treated mice.

## Conclusions

This study has highlighted the capabilities and potential of MSI to enhance investigation of the MGB axis through the detection and discovery of molecules involved in MGB communication. Here we show that, despite significant changes in gut microbiota, neurotransmitters are not significantly changed in the brain. As the significance of the MGB axis is still being realised, MSI offers a unique opportunity to understand the complexity of these interactions by identifying both the known and unknown mediators of host-microbe communication.

## Declarations

### Ethics approval and consent to participate

Approval for animal procedures was given prior to their initiation by internal University of

Manchester and University of Glasgow ethics committees and by the U.K. Home Office under licenses 70/7815, PPL40/4500, P64BCA712 and P78DD6240.

### Consent for publication

Not applicable

### Availability of data and material

The datasets during and/or analysed during the current study available from the corresponding author on reasonable request.

### Authors’ contributions

H.H., L.M.M., R.J.A.G., R.B. and D.M.W. planned the studies. H.H., L.M.M., N.S., J.S., G.H. R.A.B., S.L.B., A.S.M., and D.M.W. conducted experiments. H.H., L.M.M., N.S., J.S., G.H., S.M., R.J.A.G., R.B. and D.M.W interpreted the studies. H.H., L.M.M., R.J.A.G., R.B. and D.M.W. wrote the first draft of the paper. All authors reviewed, edited and approved the paper. A.S.M., S.M., R.J.A.G., R.B. and D.M.W. obtained funding.

## SUPPLEMENTARY INFORMATION

### Supplementary methods

#### MALDI-MSI analysis

For quantification analysis of HMG, MALDI-MSI analysis was performed using the following parameters: α-Cyano-4-hydroxycinnamic acid (CHCA) matrix was applied at 5 mg/mL concentration in 50% acetonitrile, 50% water with 0.1% trifluoracetic acid using an automated matrix applicator (HTX technologies, Chapel Hill, NC, US); 8 passes were performed using a nozzle temperature of 75°C, velocity of 1100 mm/min, flow rate of 80 μL/min, and gas pressure of 6 psi. 9-aminoacridine matrix was used at a concentration of 10 mg/mL in 80% methanol 20% water and sonicated for 20 min. The matrix was applied using an automated matrix applicator (HTX technologies, Chapel Hill, NC, US) for 3 passes, using a nozzle temperature of 75°C, gas pressure of 6 psi, velocity of 1120 mm/min and flow rate of 80 μL/min.

MALDI-MSI was carried out on a Rapiflex MALDI-TOF instrument (Bruker Daltonics, Bremen, Germany) with a 10 kHz smartbeam laser. A spatial resolution of 50 μm was achieved using a single 5 μm laser beam mode, with 600 laser shot over an area of 50×50 μm. Results were analysed using Fleximaging v5 software and the data was normalized to total ion count.

#### 3-hydroxy-3-methylglutaric acid (HMG) and pantothenic acid (vitamin B5) quantitation

Labelled standards for 3-hydroxy-3-methyl-d3-pentanedioic acid and vitamin B5 (*di-β-alanine*-^13^C_6_,^15^N_2_) calcium salt (Sigma-Aldrich, Poole, Dorset, UK) were spotted onto a brain section using a Preddator reagent dispenser (Redd & Whyte, London, UK). Brain sections with the spotted standards were imaged alongside the experimental brain sections from SPF and GF mice using DESI-MSI for vitamin B5 and MALDI-MSI for HMG. The standards were then used to generate standard curves for mean intensity against concentration for each standard spot. These values were used to calculate the concentration of the endogenous molecule across the experimental brain sections considering the thickness and area of tissue section analysed.

## Supplementary figures

**Figure S1.**
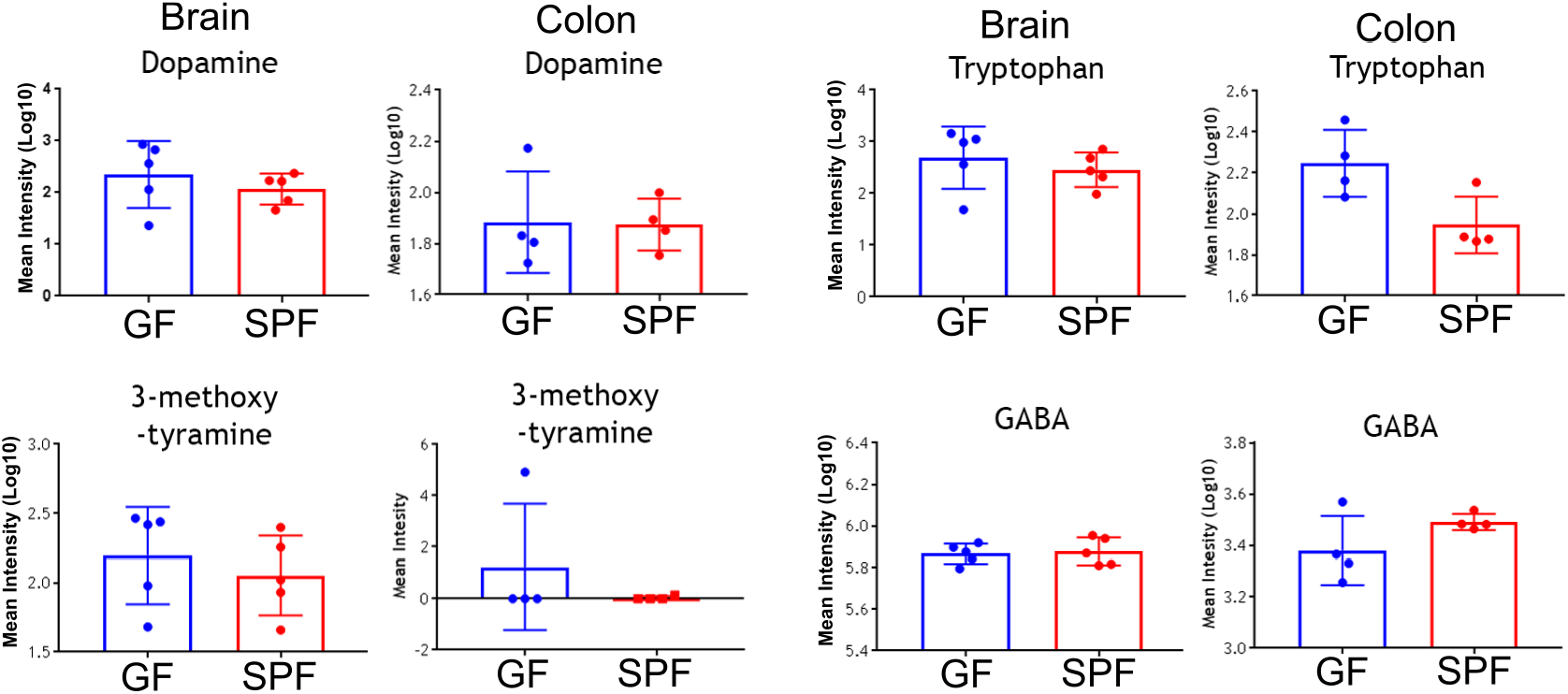
Effects of the gut microbiome on neurotransmitters, neurotransmitter precursors and neurotransmitter metabolites in the murine brain and gut. Bar plots showing the average metabolite abundances in the brain and colon of germ-free mice (GF) compared to specific-pathogen-free mice (SPF). Error bars represent standard deviation. Brain N=5, colons N=4.

**Figure S2.**
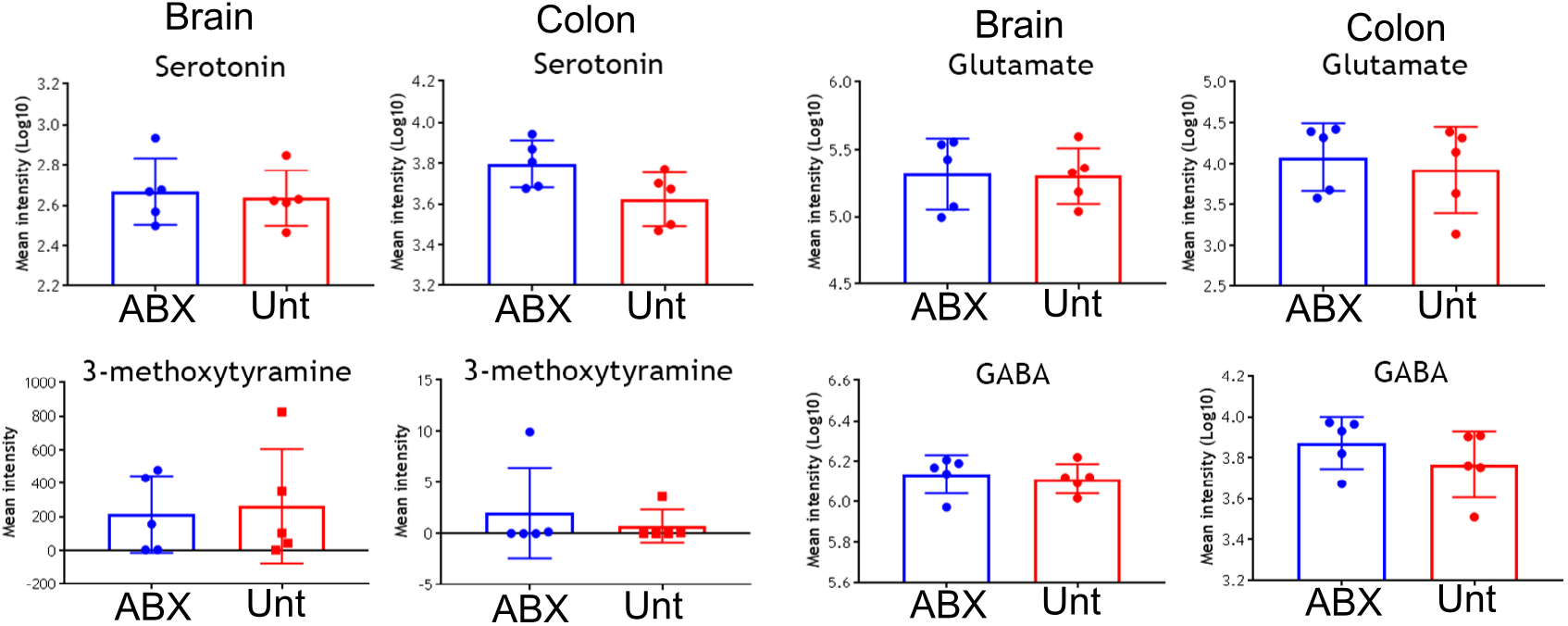
Effects of the gut microbiome on neurotransmitters, neurotransmitter precursors and neurotransmitter metabolites in the murine brain and gut. Bar plots showing the average metabolite abundances in the brain and colon of one week antibiotic treated mice (ABX) compared to untreated control mice (Unt). Error bars represent standard deviation. N=5.

**Figure S3.**
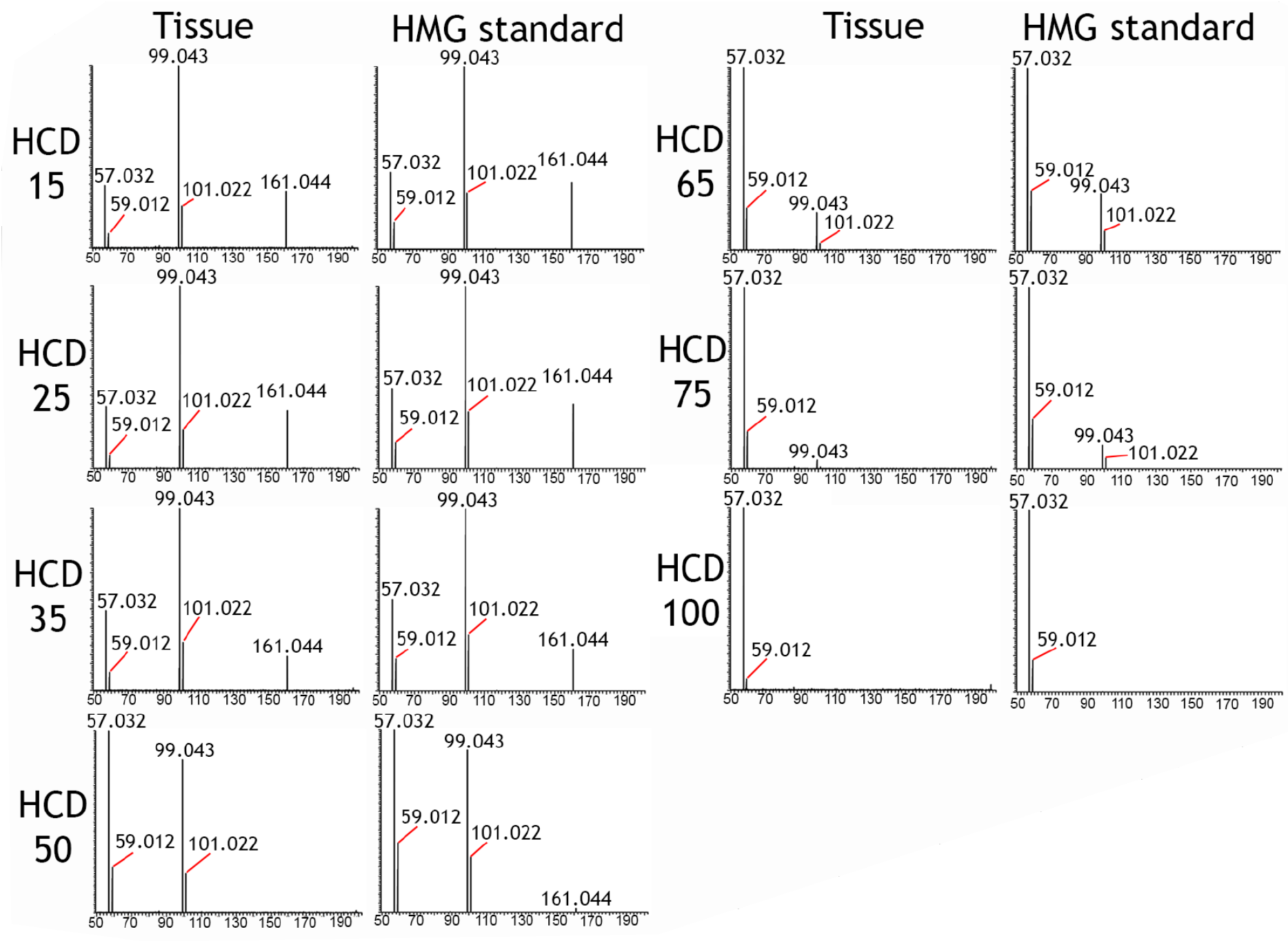
Identification of metabolite at *m/z* 161.0446. Product ion spectra from MS/MS analysis of the metabolite at m/z 161.0446 compared to product ion spectra from MS/MS analysis of 3-hydroxy-3-methylglutaric acid standard. The spectra show the m/z range across the x-axis and arbitrary unit on the y-axis. The MS/MS analysis was performed at different high-energy collision dissociation voltages (HCD 15-100).

**Figure S4.**
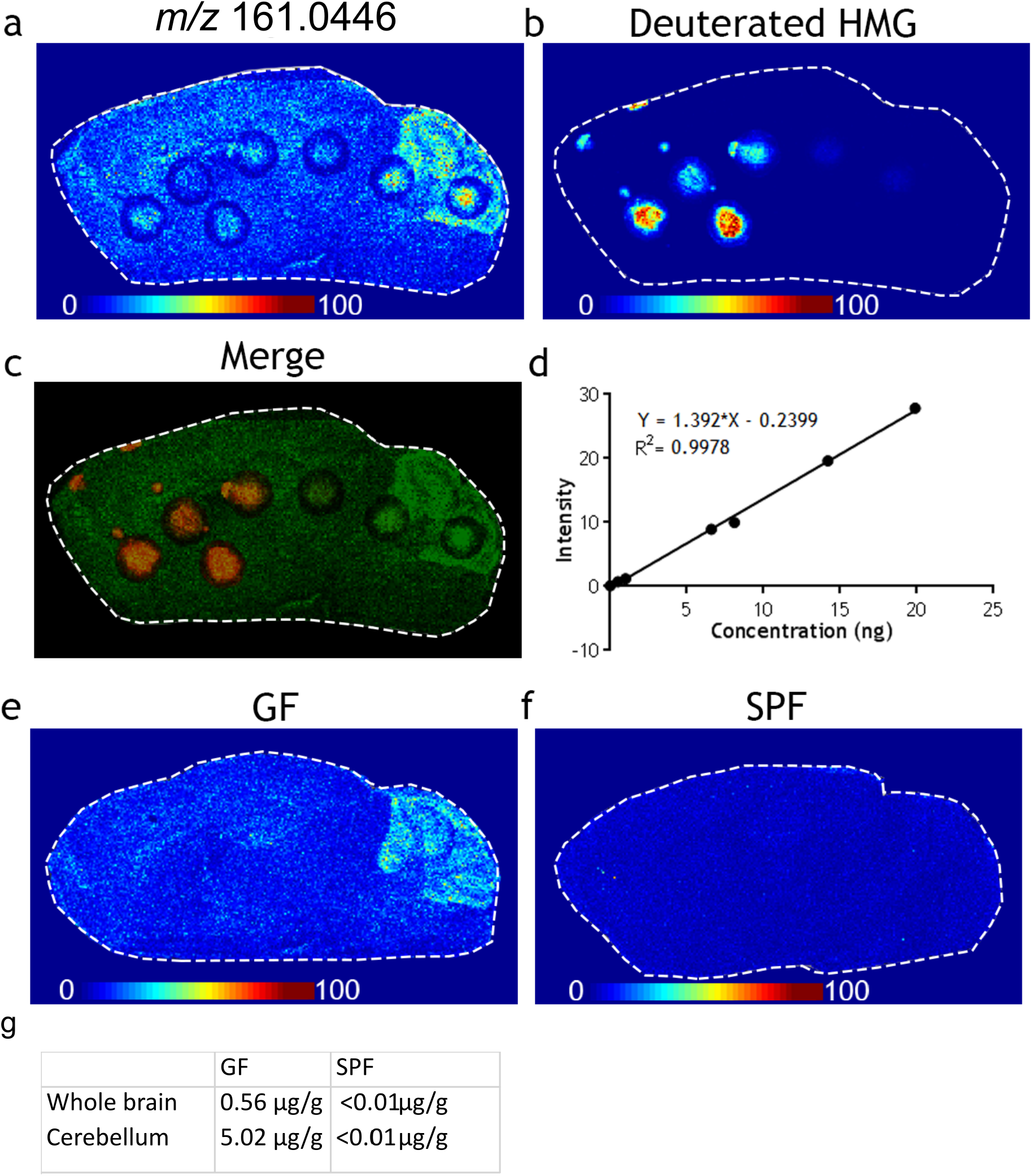
Absolute quantification of HMG in GF and SPF brain. **(a)** MALDI-MSI ion image of HMG (*m/z* 161.044) in the GF brain, which has been spotted with various concentrations of deuterated HMG standard, shown in (b). (c) Overlay of endogenous HMG and deuterated HMG standard. (d) Calibration curve obtained from the various concentrations of deuterated HMG standard spotted on the brain section. (e-g) Comparison of the HMG MALDI-MSI ion images in the GF and SPF mouse brain and absolute quantification values across the whole brain and the cerebellum.

**Figure S5.**
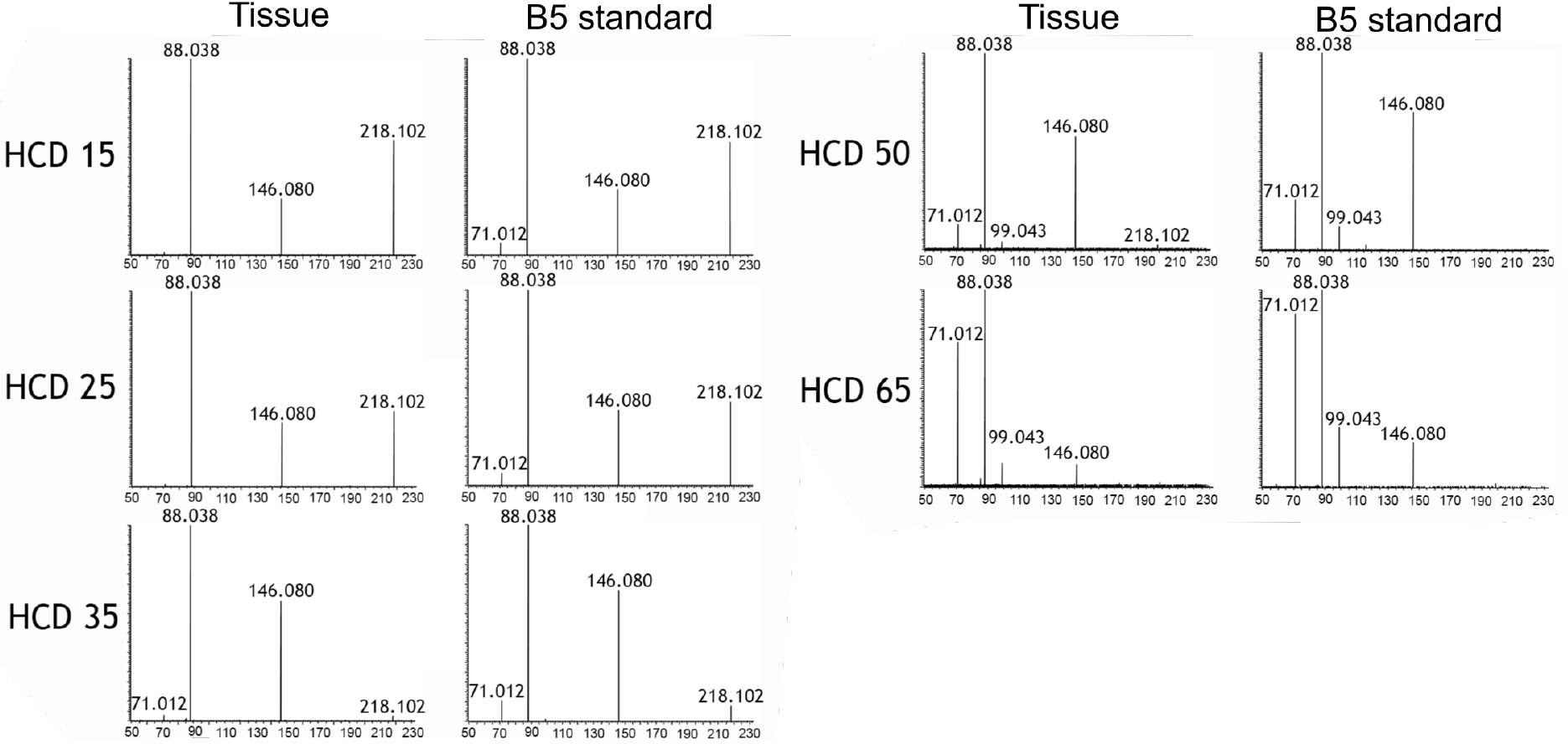
Identification of the metabolite at *m/z* 218.102. Product ion spectra from MS/MS analysis of the metabolite at m/z 218.102 compared to product ion spectra from MS/MS analysis of vitamin B5 standard. The spectra show the m/z range across the x-axis and arbitrary unit on the y-axis. The MS/MS analysis was performed at different high-energy collision dissociation voltages (HCD 15-65).

**Figure S6.**
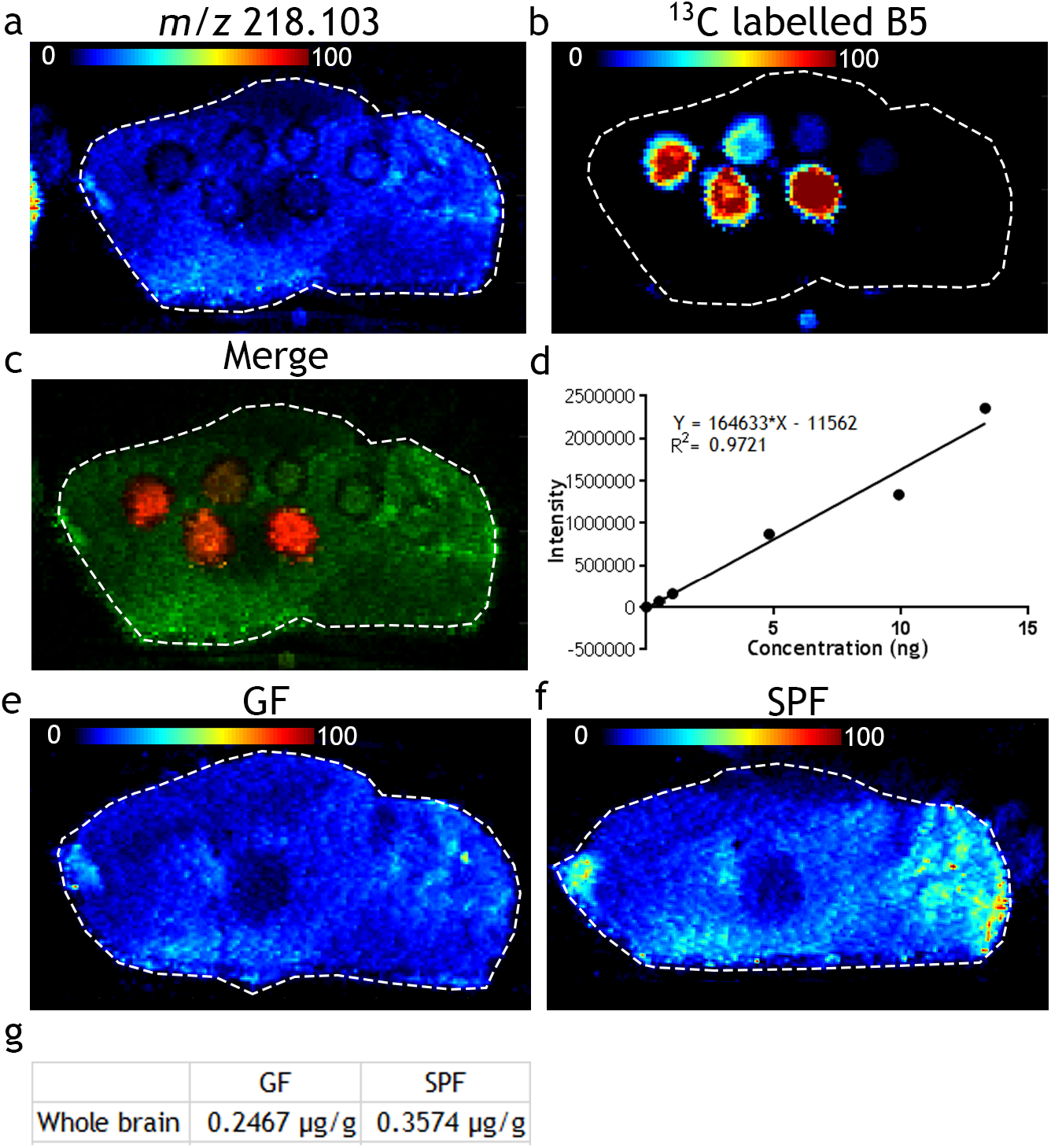
Absolute quantification of vitamin B5 in GF and SPF brain. **(a)** DESI-MSI ion image of B5 (*m/z* 218.103) in the GF brain, which has been spotted with various concentrations of ^13^C labelled B5 standard, shown in (b). (c) Overlay of endogenous HMG and ^13^C labelled B5 standard. (d) Calibration curve obtained from the various concentrations of deuterated ^13^C labelled B5 standard spotted on the brain section. (e-g) Comparison of the B5 DESI-MSI ion images in the GF and SPF mouse brain and absolute quantification values across the whole brain and the cerebellum.

